# Male mice are particularly vulnerable to cognitive impairment following mTBI

**DOI:** 10.64898/2026.02.21.707169

**Authors:** James T. Neal, Aimee Bertolli, Georgina M Aldridge, Eric B. Emmons

## Abstract

Traumatic brain injuries (TBIs) result from impact to or rapid displacement of the brain and can lead to various neurological deficits involving working memory, decision-making, and anxiety. While large-scale effects of brain damage are well-described for more severe TBIs, less is known about the extent and duration of cognitive deficits at the mild level. Interval timing can provide a helpful window into cognition in mice and humans. Interval-timing behavior is impaired in a wide range of neuropsychiatric disease states, such as Parkinson’s disease. Furthermore, novel object recognition (NOR) and the Barnes maze (BM) tests are valuable assays for evaluating spatial learning, working memory, and anxiety-like behavior in mice. Here, we employed a weight-drop model of mild TBI (mTBI) to investigate changes in internal cognitive states resulting from mTBI treatment. mTBI mice were not significantly impaired in either interval timing or NOR, but they demonstrated impaired spatial memory in the Barnes Maze. Interestingly, within-sex comparisons revealed impairments in male mTBI mice in the interval-timing task and the NOR, suggesting that male and female mice may be differently affected by mTBIs.

## Introduction

Common in falls, motor vehicle accidents, and contact sports, a mild traumatic brain injury (mTBI) is the most common subtype of TBI, accounting for about 75% of all TBIs (Blaylock & Maroon, 2011; *Multiple Cause of Death Data on CDC WONDER*, n.d.). It is estimated that around 0.6 percent of the global population experiences a mTBI annually, equating to about 42 million people worldwide, with many of these going undiagnosed (Gardner & Yaffe, 2015).

Traumatic brain injuries are characterized by the DSM-5 as rapid displacement of the brain within the skull caused by an impact or a fast acceleration/deceleration (*Diagnostic and Statistical Manual of Mental Disorders, Fifth Edition (DSM-5)*, n.d.). Although specific diagnostic criteria are variable, most criteria for a mTBI require loss of consciousness lasting less than 30 minutes, post-traumatic amnesia lasting no longer than 24 hrs, and no evidence of trauma upon neuroimaging (Katz et al., 2015). Often accompanying a mTBI is post-concussion syndrome, a cluster of symptoms such as headache, dizziness, irritability, fatigue, and memory impairment (Blaylock & Maroon, 2011; Katz et al., 2015). Although these symptoms are typically transient, repetitive models of mTBI in rodents, as well as studies of chronic traumatic encephalopathy, have demonstrated more persistent cognitive impairment, implying that the pathophysiological effects of each mTBI may be cumulative and make it difficult to fully recover (Blaylock & Maroon, 2011; Stern et al., 2011; Wang et al., 2019).

The mechanism of injury for a TBI can be divided into two parts: the primary and secondary injury (Gardner & Yaffe, 2015). The primary injury results from the immediate mechanical force against the brain, often causing hemorrhage, axonal shearing, and temporary disruption of the blood-brain barrier (Blaylock & Maroon, 2011; Liu et al., 2022). The secondary injury is caused by post-injury molecular cascades. It lasts much longer, impairing some aspects of basic cognition such as spatial memory and anxiety for up to six months in some rodent models (Boyko et al., 2022; Luo et al., 2014, 2017). The mechanisms by which mTBIs induce these cognitive deficits are still relatively unclear; however, studies in mouse models have identified some biomarkers of neurodegeneration, such as hyperphosphorylated tau, microglial activation, astrocytosis, and neurofilament light protein as correlates of mTBI (Acosta et al., 2013; Blaylock & Maroon, 2011; Hernandez-Ontiveros et al., 2013; Kane et al., 2012; Ojo et al., 2013; Petraglia et al., 2014; Stern et al., 2011; Wang et al., 2019).

Time perception is one aspect of basic cognition that is impaired in many neuropsychiatric disease states, such as Parkinson’s disease, schizophrenia, and ADHD (Singh et al., 2023; Vaidya & Stollstorff, 2008; Yoon et al., 2008). The prefrontal cortex and striatum are critical regions in temporal processing, and impairments in these regions result in time-related deficiencies (Emmons et al., 2017; Mello et al., 2015; Merchant & Averbeck, 2017; Narayanan et al., 2012). Interval timing is an effective method of assaying cognition involving these critical brain regions and is a method seldom applied to models of TBI (Scott & Vonder Haar, 2019). Here, we used the “switch task” variant of interval timing, an operant-conditioning task designed to measure a rodent’s response to time-based cues (Balci et al., 2008; Larson et al., 2022; Stutt et al., 2024). The interval-timing switch task is able to assess the animal’s knowledge of two different time intervals, and, importantly, when they choose to make the “switch” from one nosepoke response port to the other.

The Barnes maze is another assay used to measure basic cognition, relying on the assumption that a rodent should be able to learn and remember the location of an escape from a potentially perilous environment (Pitts, 2018). Consisting of an elevated platform with various false escape holes and one real escape hole, the Barnes maze measures spatial learning, memory, and cognitive control (Gawel et al., 2019). As training progresses, the mouse learns to pick up on spatial cues to find and remember the location of the escape port. A novel addition to the Barnes maze is the Spatial Training and Rapid Reversal protocol (STARR protocol; Bertolli et al., 2024). The STARR protocol introduces barriers similar to a radial arm maze, forcing mice to utilize different search strategies than what is possible in a traditional Barnes maze. Additionally, during the STARR protocol, the location of the escape is abruptly changed, measuring the animal’s cognitive flexibility as it is forced to adapt to unexpected circumstances.

Here, we sought to explore sex-related similarities and differences in various executive function tasks following mTBI. Because a large portion of our understanding of executive function tasks derives from studies using only male rodents, it is important to further investigate behavioral sex differences that may naturally exist in such assays. In research that has explicitly compared sex as a variable in spatial learning and memory tasks, some studies have found mild differences between sexes, while some studies have found no difference at all (Stevanovic et al., 2022; Tsao et al., 2023). Additionally, mice exhibit sex-specific sensitivities to different amounts of handling–indeed, minimally habituated female mice exhibit a decreased investigation of objects in a spatial object task (Stevanovic et al., 2022). This result was not seen in males and was attenuated by increased handling time prior to experimentation.

In this experiment, we hypothesized that induction of a mild TBI would result in short-term, widespread cognitive impairment involving deficits in memory, interval-timing behavior, and cognitive flexibility in both male and female mice. We sought to a) characterize the extent of cognitive impairment induced by a single mTBI event in male and female mice and b) to explore the timeframe in which these deficits occur. mTBI mice were globally impaired in the STARR protocol component of the Barnes maze, with no consistent deficits observed in the switch interval-timing task or the novel object recognition task. However, when sex was considered as a variable, male mice were more susceptible to the mTBI treatment, with disrupted performance in the switch interval-timing task and the novel object recognition task. This finding suggests that there may be sex-specific cognitive vulnerabilities to even mild traumatic brain injury.

## Materials and Methods

### Animal Subjects

20 three-month-old C57BL/6J mice were divided evenly into two groups, a “sham” control group and a mTBI group, each group consisting of five males and five female mice. One group was induced with a mTBI following training in a switch interval-timing operant conditioning task, and the other group served as the control. Mice were housed in cages of two or three with unlimited access to water and were food-deprived until they reached about 80% of their starting weight to increase motivation in the operant conditioning paradigm (Lewon & Hayes, 2015). All procedures were approved by the University of Iowa IACUC and performed according to their guidelines.

### Interval-timing switch task

Interval timing was measured by an operant-conditioning paradigm known as the “switch task.” The switch task is designed to measure the subject’s estimation of a time interval based on their responses to time-based cues (Balci et al., 2008). It consists of three shaping stages that progressively approach the final version of the task.

***Stage 1.*** In the first stage, mice learn to associate the ports on the front wall with the activation of a sucrose pellet dispenser on the back wall. Once a mouse reached 10 rewards in a 90-minute session, they were advanced to the next stage.

***Stage 2.*** In the second stage, a trial could be initiated with a nosepoke to the port on the back of the operant-conditioning chamber. Once initiated, the mouse would begin either a 6-second or 18-second trial at random. A 6-second trial would reward the mouse for the first nosepoke to the port designated as the short-trial port (left or right port, counterbalanced across all of the mice) after 6 seconds had passed. An 18-second trial would reward the mouse for the first nosepoke to the port designated as the long trial port after 18 seconds had passed (Figure 1). In this stage, mice learned to associate one port with the fastest potential reward and the other port with a delayed reward. Intertrial intervals were randomized around an average of 90 seconds to ensure that mice did not learn to time the intertrial interval. Mice were advanced to the next stage after four or five days.

**Figure 1.**
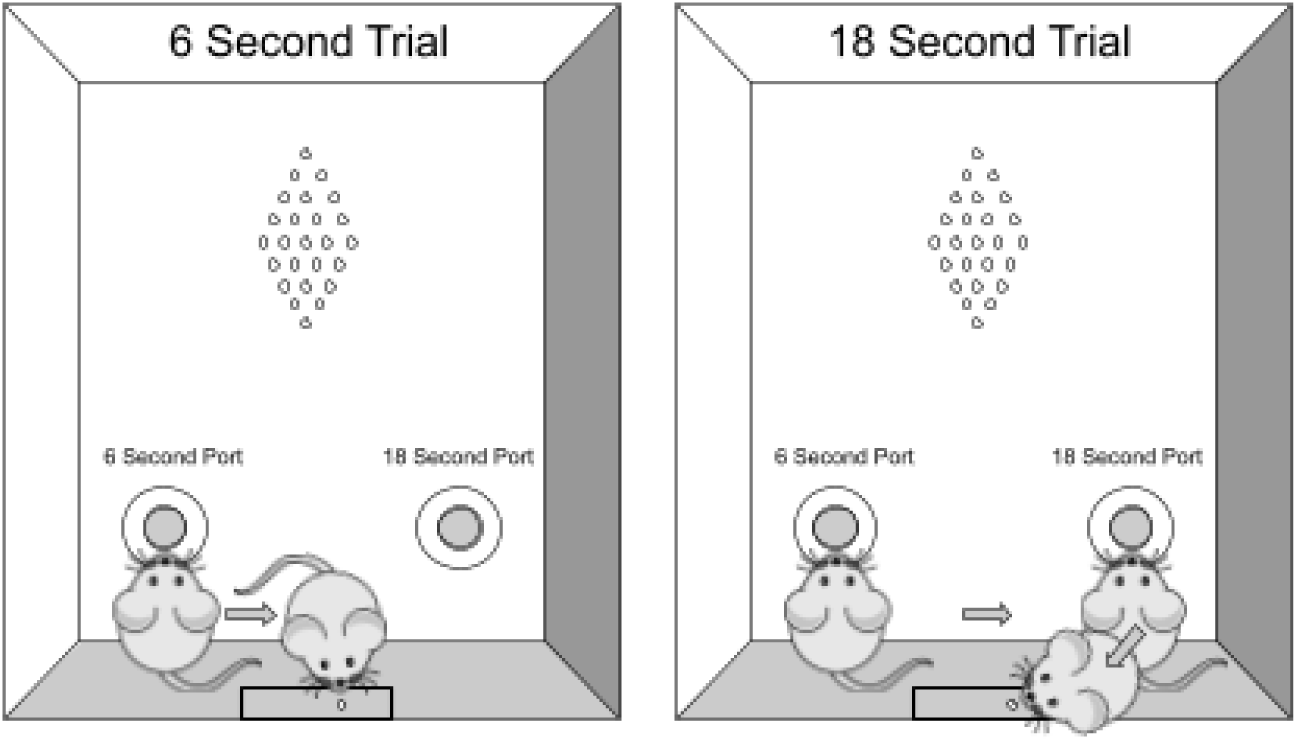
Operant Conditioning Box Layout During a 6-Second and 18-Seconf Trial. The mouse initiates a trial with a nosepoke to a port on the back wall (not depicted here).

***Stage 3.*** The third stage required mice to visit the short trial port before entering the long trial port during an 18-second trial. This reinforced that each trial would start (6 or 18 seconds) with nosepoke response at the short-trial port, only switching to the long-trial port when they perceived that 6 seconds had passed. The time at which the mouse perceived the trial to be an 18-second trial is known as the “switch time” and is what was used to estimate changes in interval timing. The switch time was defined as the point at which the mouse removed its head from the infrared sensor in the short-trial port. Mice ran the switch interval-timing task for 90 minutes each day.

Mice were trained in Stage 3 of the task for at least 16 days prior to mTBI induction (or sham treatment if they were a control group mouse). All mice continued the switch interval-timing task for 12 days following their treatment. Data was analyzed for only 18-second trials where the following was measured: switch time, the coefficient of variance (CV) of switch time, and the ratio of trials correct. A correct trial was considered any 18-second trial in which the mouse visited the short-trial port before switching to the long-trial port to receive a reward. The short-trial port was assigned as the left port for half of the mice, while in the other half, it was assigned as the right port. This was done in order to avoid a bias for one of the ports.

### Weight-Drop Method

mTBIs were induced via the weight-drop method, which is a non-invasive, closed-head model of diffuse mTBI (Kane et al., 2012). Before mTBI induction, all mice were placed in a chamber where they were anesthetized with 5% isoflurane gas until unresponsive. The mice were then positioned onto a 2.5 cm-thick sponge pad with a 0.5-cm layer removed to allow for consistent placement of the mouse’s body and head. The sponge pad was used to allow for subtle movement of the brain within the skull to model common TBIs occurring in sports and vehicle accidents (Kane et al., 2012; Wang et al., 2019). A 1-cm diameter fiberglass tube was positioned so that the opening was evenly spaced between the ears and just posterior to the eyes. A 21-g weight of 1-cm diameter was dropped from a height of 50 cm onto the mTBI group (Wang et al., 2019; Yang et al., 2013). This height was adapted from other similar approaches to induce a minimally severe mTBI in C57BL/6J mice (Shishido et al., 2019; Wang et al., 2019).

### Novel object recognition

The novel object recognition (NOR) task was used to measure anxiety-like behavior and working memory (Antunes & Biala, 2012; Lueptow, 2017; O’Shaughnessy et al., 2023). Mice first underwent an open field test to assess movement in an empty chamber, followed by three novel object trials according to the sequence in Figure 2. Trial 1 consisted of two of the same objects in the chamber (both of which were novel at that stage). Trials 2 and 3 both had one novel and one familiar object. Visits to and time spent at the novel and familiar objects were recorded, as well as total distance traveled within the NOR chamber. Objects were designed and 3D-printed in-house using PLA filament. The three object types were designed and tested such that they were the same weight, height, and had shapes that were difficult for the mice to climb onto. The color and name combos–names roughly corresponding to their shape–were purple blob, golden trophy, and gray spike. The location and starting object (which would become the familiar object for Trials 2 and 3) were counterbalanced across male and female mice as well as the sham and mTBI groups.

**Figure 2.**
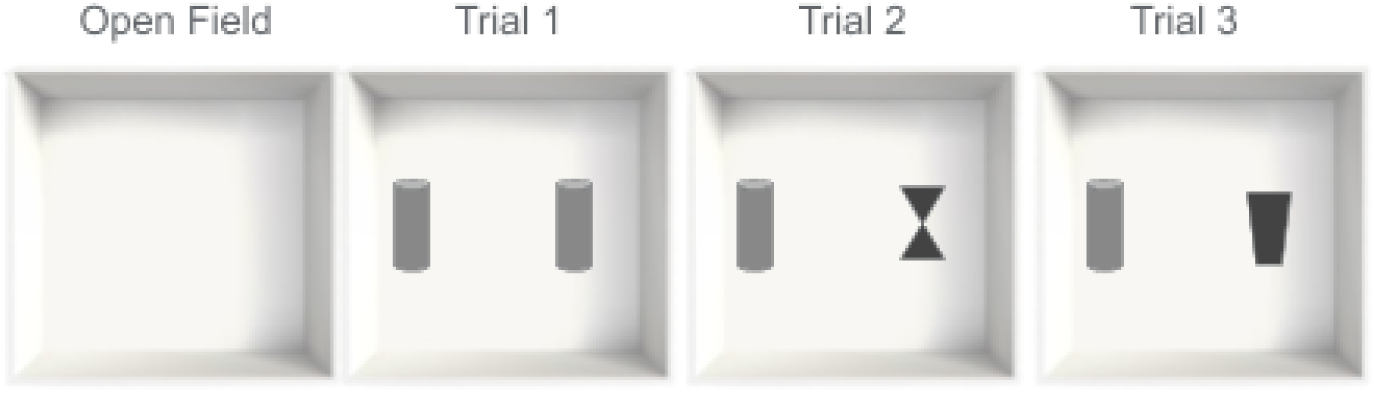
Sequence of Events in the Novel Object Recognition Protocol. Open field begins with no objects. Trial 1 has two novel objects. Trials 2 and 3 both have one familiar and one novel object.

### Barnes Maze

Spatial learning and working memory were assessed with a modified form of the Barnes maze, building on standard versions with the addition of a fourth day of behavior–the STARR protocol, described below–one week following the initial three days (Bertolli et al., 2024). The maze used was an elevated, circular platform with a 48-inch diameter and 10 evenly-spaced holes along the circumference, as depicted in Figure 3. On a given trial, one of the ten holes, the target, leads to an escape shuttle, while the rest are obstructed by a plastic grate. A trial was initiated by placing the mouse in the center of the platform. Once the mouse found the target shuttle, the trial was concluded. A tunnel was then connected, allowing the mouse to self-exit the shuttle to its own cage. The maze was placed in a well-lit room with visual cues along each wall, allowing the mouse to use these cues to navigate the spatial environment (Bertolli et al., 2024). Four days after treatment, mice were trained for three consecutive days on the same target in the same location of the maze. Day 1 began with a habituation trial, after which there were five training trials using the target. Days 2 and 3 both began with a 90-second probe trial, where all holes were sealed, and the mouse’s activity was recorded. On Day 2, this was followed by six training trials. On Day 3, the six training trials were followed by a 3-minute probe trial. One week following Day 3, we conducted a new modification of the Barnes maze developed by our collaborators termed the STARR (<u>S</u>patial <u>T</u>raining <u>A</u>nd <u>R</u>apid <u>R</u>eversal) protocol (Bertolli et al., 2024). This additional day presents the mice with additional contingencies and two new targets, allowing for the assessment of working memory and behavioral flexibility. In the STARR protocol, acrylic dividers (16” x 3.5” made of 0.25” clear acrylic; Professional Plastics, Sacramento CA) are positioned to extend 2/3 of the way into the maze, forcing the mice to be more decisive in their exploration of the maze instead of using a serial exploration strategy. On Day 4 of the STARR protocol, the first trial is a probe trial of memory retention from the previous week’s location and is run in standard form without barriers. Trials 2 and 6 have a new location, distinct from the location learned over the original three days of training. During trials 2-5, the mice acclimate to the acrylic dividers and try to rapidly acquire a new spatial location. Trials 6-9 again switch the location of the escape port, measuring the mouse’s cognitive flexibility or tendency towards perseveration. Trial 10 is a final probe trial of the STARR protocol–the mouse must either visit all 10 arms or spend up to 10 minutes exploring. The program ANY-Maze was used to score Barnes maze and NOR data (ANY-Maze, Stoelting Co., Wood Dale, IL). ANY-Maze tracked the number of times the mouse investigated a hole (entries) and the number of new holes it investigated (unique entries). A Pearson correlation between ANY-Maze-scored entries and hand-scored entries found a very strong positive correlation, suggesting that the ANY-Maze automated tracking was highly accurate (Figure 4). There are several different outcome measures that can be assessed throughout the four days of the Barnes maze and STARR protocol–for a comprehensive overview, see Bertolli et al. 2024.

**Figure 3.**
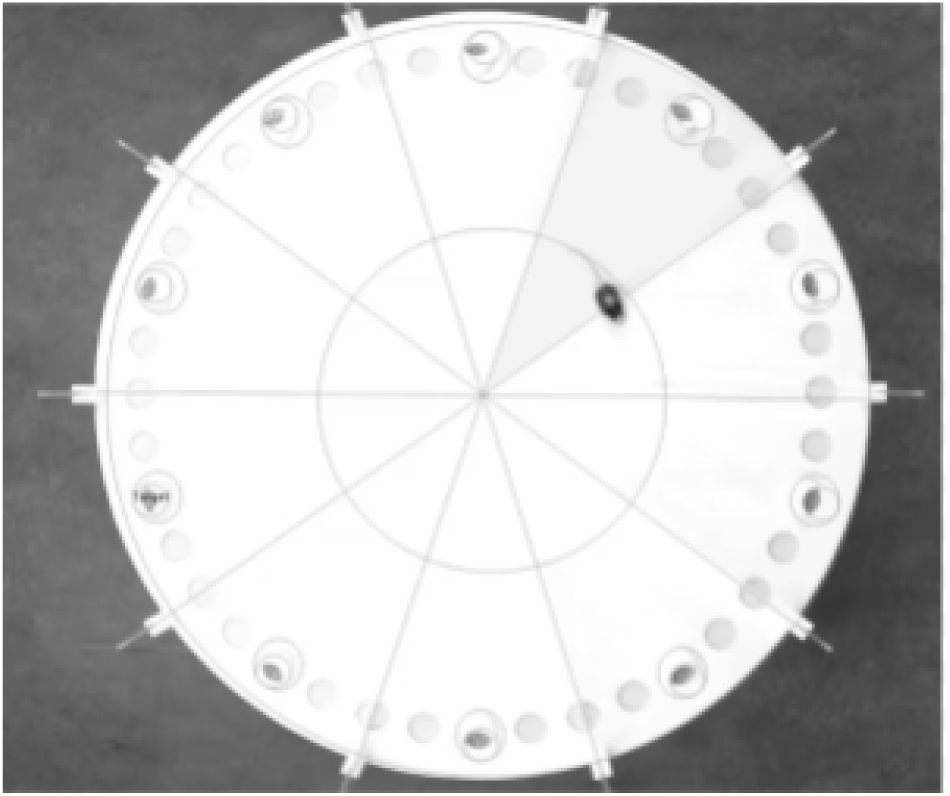
Aerial Image of the Barnes Maze Being Scored by AnyMaze. The image depicts the maze on Day 4, after acrylic dividers have been introduced. Entries, unique entries, and working memory errors were scored based on when the mouse’s nose enters the circle outlining the hole on the maze.

**Figure 4.**
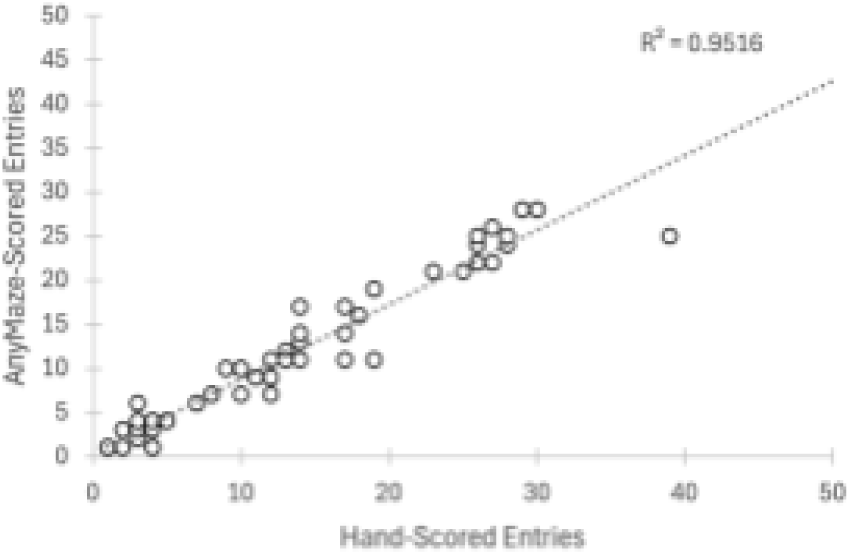
Anymaze-Scored Entries vs. Hand-Scored entries. To verify the accuracy of the video tracking program AnyMaze, a linear regression of AnyMaze-scored entries and head-scored entries was performed R^2^ = 0.9516, indicating a high correlation.

### Statistics

All statistical tests were conducted in R. Unpaired t-tests were calculated using: t.test(data ∼ Condition, alternative = “two.sided”, paired = FALSE)

In the interval-timing task, variability in switch times was calculated using a coefficient of variance (CV). Post-injury switch time CVs, ratios of trials correct, and switch times were normalized and expressed as a percent of the respective pre-injury baseline data to account for individual variability. A normalized CV of 100 percent indicated no difference from pre-injury data. Variables in the interval-timing task were compared between sham and mTBI groups with unpaired, two-tailed t-tests.

Unpaired two-tailed t-tests were performed to compare dependent variables between sham and mTBI mice in the novel object recognition (NOR) task. Normalized discrimination between novel and familiar objects in the NOR test was calculated using the protocol outlined by Lueptow et al. The time spent at the familiar object was subtracted from the time spent at the novel object and divided by the total time exploring either object, outputting a value between -1.0 and 1.0 (Lueptow, 2017). A value closer to 1.0 indicates a preference toward the novel object, and a value closer to -1.0 indicates a preference toward the familiar object.

As preliminary analyses did not indicate a significant difference between male and female mice in the Barnes maze, group comparisons were collapsed across sex. For Unique Visits on Day 4 of the STARR Protocol, mice that have deficient memory of a 10-hole Barnes maze will explore 5.5 or more holes before finding the target–a one-sample Wilcoxon sign-ranked test was run in R for this analysis with the following parameters:

wilcox.test(data $ Unique Entries, mu = 5.5)

All other Barnes Maze analyses utilized mTBI and sham group comparisons via unpaired, two-tailed t-tests comparing repeat entries to a given zone of the maze, unique entries across the maze, and distance traveled.

### Transparency and Openness

Experiments were conducted during a 12-week summer fellowship at the University of Iowa, which dictated our sample size of 10 male and 10 female mice. All animals are included in each analysis. Data are available on Zenodo at the following DOI: https://doi.org/10.5281/zenodo.18717441. Data were analyzed using R, version 4.4.3 (The R Foundation for Statistical Computing, 2025). This study’s design and its analysis were not pre-registered.

## Results

### Switch interval-timing task

Switch interval-timing behavior was assessed at three different post-injury timepoints: 1-4, 5-8, and 9-12 days post-injury (Figure 5). Post-treatment percentages of switch times, switch time CVs, and ratio of trials correct were compared to the 4 days prior to treatment. Treatment group alone had no effect on any dependent variable at any time point (*p > 0.05*; Figure 5A-F). However, 1-4 days post-treatment, male mTBI mice had lower percent switch times than sham males, indicating that they switched to the long-trial port earlier compared to their pretreatment performance (all results reported as mean ± standard error: 104.7% ± 1.92 vs. 93.3% ± 2.48; *p < 0.01*; Figure 5D). This effect persisted into the 5-8 day post-treatment period (99.7% ± 1.53 vs. 92.1% ± 1.56; *p < 0.05*; Figure 5E) but did not reach significance in the 9-12 day period (*p > 0.05*; Figure 5F). Female mTBI mice, however, showed no significant difference from sham females in any dependent variable at any time point.

**Figure 5.**
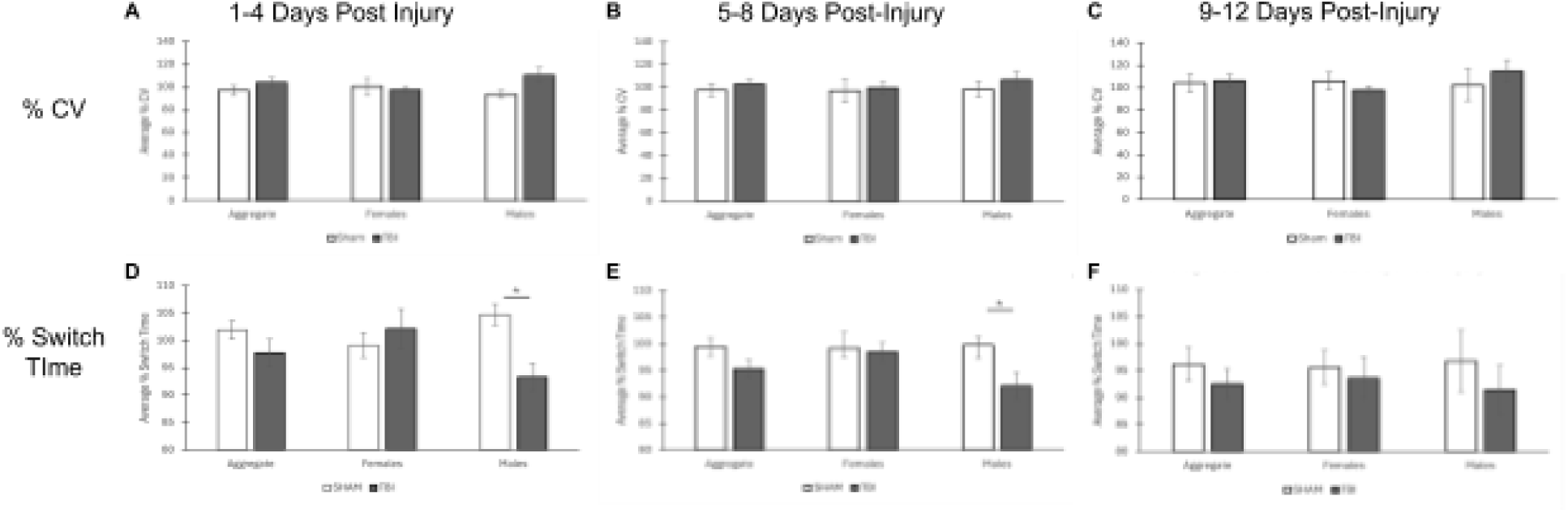
Performance in the Switch Operant Conditioning Task. **(A)** Average percent coefficient of variance 1-4 days injury. **(B)** Average percent coefficient of variance 5-8 days post injury. **(C)** Average percent of variance 9-12 days post injury. **(D)** Average percent switch time 1-4 days post injury. **(E)** Average percent switch time 5-8 days post injury. **(F)** Average percent switch time 9-12 days post injury. 100% indicates the pre-treatment baseline. A value below 100% indicates a post-treatment decrease, while a value above 100% indicates a post-treatment increase, An asterisk denotes a p < 0.05. Error bars represent the standard error.

### Novel Object Recognition

In trial 1 of the novel object recognition (NOR) task, mTBI mice made more visits to the two objects–both objects are novel on this first trial–compared to sham (sham: 117.5 s ± 6.03 vs. mTBI: 136.3 s ± 8.44; *p < 0.05*; Figure 6A). However, there was no statistical difference in the length of time spent at the novel objects, due to high variability between mice (sham: 171.45 s ± 29.98 vs. mTBI: 218.65 s ± 52.63; *p = 0.485*; Figure 6B). In trials 2 and 3–when the novel object was introduced along with the familiar object from Trial 1–mTBI mice showed no significant difference in discrimination between novel and familiar objects compared to sham, as depicted in Figure 6C (*p > 0.05*). Male mTBI mice, however, showed a preference for the familiar object as compared to the sham males’ moderate preference for the novel object (sham: 0.26 ± 0.059 vs. mTBI: -0.06 ± 0.093; *p < 0.01*; Figure 6C). The higher preference of the mTBI males toward the familiar object underlines a non-random association between novel and familiar objects and denotes working memory of the object presented in the previous trial. The preference toward the familiar object may be indicative of an anxiety-like phenotype. Female sham mice did not show a significant preference for either object (0.007 ± 0.090) and were not significantly different from mTBI females (0.019 ± 0.119; *p = 0.86*). Notably, while male sham mice showed a moderate preference towards the novel object, female sham mice showed almost no preference for one object over the other.

**Figure 6.**
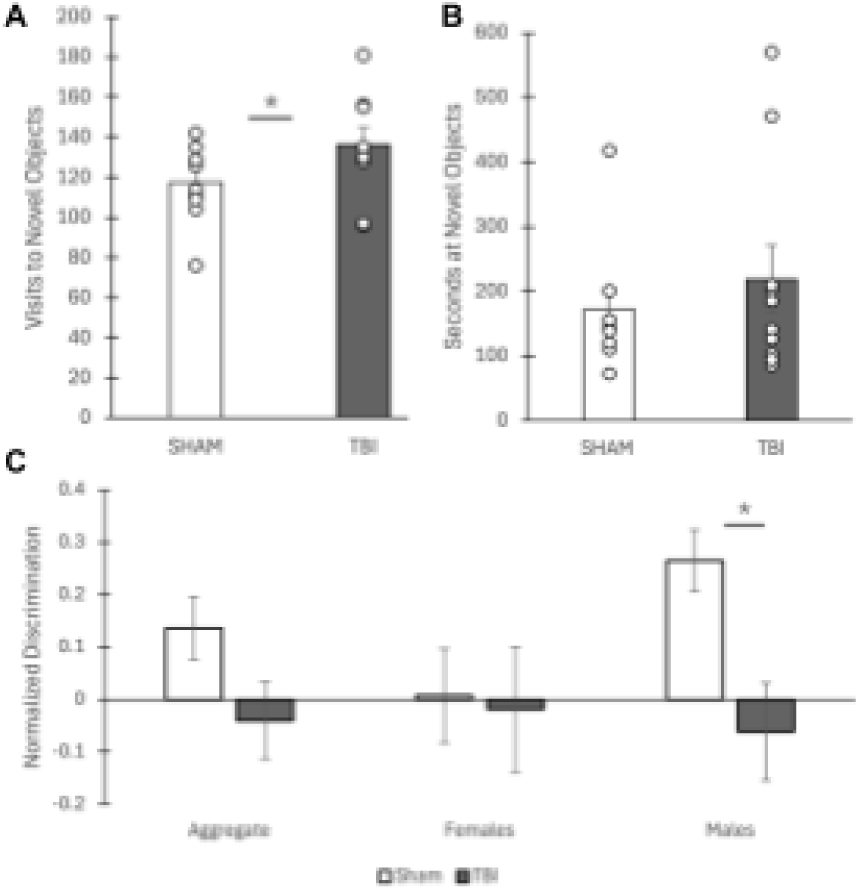
Performance in the Novel Object Recognition Task. **(A)** Visits to either novel object in trial one of NOR. **(B)** Seconds spent at either novel object in trial one of NOR. **(C)** Normalized object discrimination in trials with a novel and familiar object. A value closer to 1 indicates a preference towards the novel object. A value closer to −1 indicates a preference towards the familiar object. An asterisk indicates p < 0.05. Error bars represent the standard error.

### Barnes Maze

No consistent differences in acquisition or in performance on the probe trials were observed for Days 1-3 of the Barnes Maze protocol. However, mTBI mice performed significantly worse than sham mice in the Day 4 STARR protocol of the Barnes Maze. This was evident in that mTBI mice explored more of the maze before finding the target, represented by a higher number of unique entries (sham: 3.68 ± 0.322 vs. mTBI: 4.83 ± 0.368; *p < 0.05*; Figure 7A). Additionally, mice with no memory of the cue location should make 5.5 or more unique visits for a 10-hole maze–sham mice were significantly below this threshold while mTBI mice were not (Wilcoxon signed rank test; sham: *p < 1 x 10^-5^*; mTBI: *p > 0.05*). mTBI mice also made more working memory errors (repeat entries) per trial on Day 4 of the Barnes Maze compared to sham mice (sham: 1.35 ± 0.398 vs. mTBI: 2.70 ± 0.523; *p < 0.05*; Figure 7B). Additionally, mTBI mice explored the maze more inefficiently, as evidenced by the distance traveled before finding the escape port (sham: 7.23 ± 1.039 m vs. mTBI: 11.95 ± 2.043 m *p < 0.05*; Figure 7C). For statistical analyses in Figure 7 A-C, Trials 3-5 and Trials 7-9 were combined together to measure behavioral flexibility following the changing contingencies of the STARR protocol (Trial 2 is when the barriers are introduced, and Trial 6 is a Reversal trial–6R on Figure 7D; Bertolli et al., 2024). No intra-sex comparison between mTBI and sham females reached significance.

**Figure 7.**
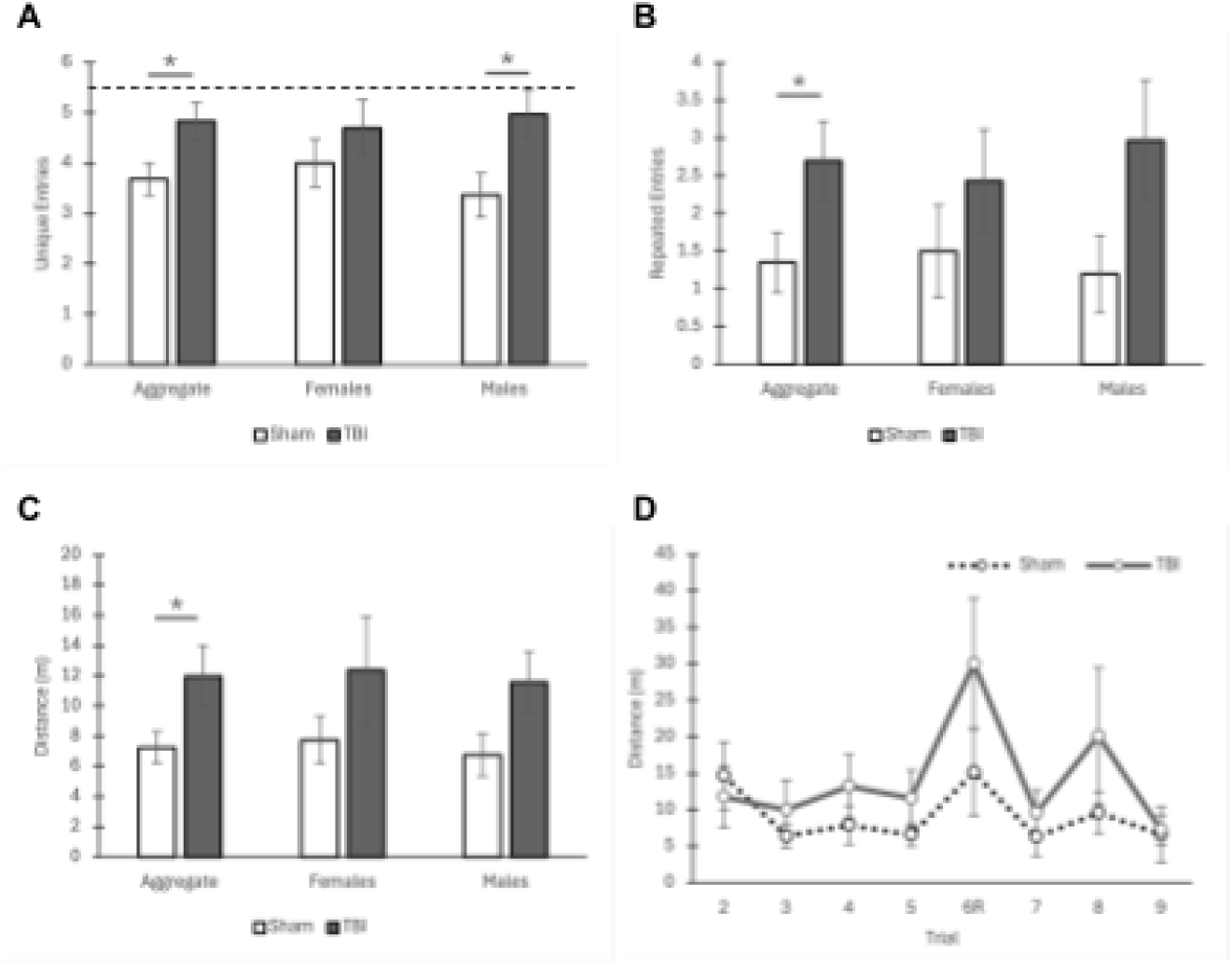
Performance in the Barnes Maze on the Day 4 STARR Protocol. **(A)** Average unique entries per trial in trials 3-5 and 7-9. The dashed line at 5.5 unique entries represents chance performance-an animal performing at or above this level demonstrates impaired memory on the task. **(B)** Average repeated entries (entries into a previously entered hole) per trial in trials 3.-5 and 7-9. **(C)** Average distance traveled per trial in trials 3-5 and 7-9. **(D)** Average distance traveled among all non-probe trials. Trial 6R indicates the reversal trial, when the target escape port is abruplty switched. An asterisk denotes a p value < 0.05. Error bars represent the standard error.

## Discussion

Here, we found evidence of subtle, sex-specific behavioral impairments following exposure to a single mTBI in mice. Across different behavioral assays probing attention and spatial memory, we found that male mice were particularly susceptible to mTBI. Female mTBI mice did not show impaired behavior in either the interval-timing switch task or the novel object recognition task. In the STARR protocol of the Barnes maze, both male and female mTBI mice showed difficulty with the flexible acquisition of new spatial locations, showing increased distance to the target and an increased number of repeat entries. These findings may suggest that mTBI mice have either impaired working memory or increased perseveration, either of which could cause an increase in the number of working memory errors and inefficiencies in learning new maze conditions. These data warrant further exploration of whether there may be neuroprotective effects of female sex hormones on the behavioral consequences of mild traumatic brain injury and concussion.

Examining the interval-timing switch task at various post-injury timepoints allowed us to estimate a timeline for cognitive impairment in male mice following a single mTBI event. Switch interval-timing behavior was shifted significantly earlier for male mTBI mice as compared to male sham mice for the first eight days following the weight-drop injury. This shift in timing behavior subsided after eight days post-injury, indicating possible recovery. We did not find any consistent effects of the single weight-drop injury on the variability of timing between the mTBI and sham groups in the switch interval-timing task, contrary to what other literature would suggest (Scott & Vonder Haar, 2019). This is likely because our study was conducted on a rigid timeline, allowing a maximum of 28 days of training prior to TBI induction. As a result, four of the mice still had relatively high variability in their timing behavior prior to the mTBI induction. This makes our comparison of the baseline switch interval-timing behavior to the post-injury behavior more difficult to interpret.

In trial one of the novel object recognition (NOR) task, mTBI mice seemed to be more exploratory than sham mice, making more visits to both objects. However, when presented with a novel and a familiar object (Trials 2 and 3), they showed a slight preference for the familiar object. The latter result is consistent with other murine NOR studies into TBI, suggesting a possible anxiety-like phenotype following mTBI treatment (Yang et al., 2013). Additionally, there was a significant difference between male and female sham mice when exposed to the novel and familiar objects. While sham males showed a preference toward the novel object, sham females showed almost no preference for either object. This is contradictory to other studies that suggest both male and female mice tend to exhibit a preference toward novel objects (Bettis & Jacobs, 2012; Lueptow, 2017; Stevanovic et al., 2022). This was an unexpected result that needs further study to determine its significance. Possible explanations include that mTBI may affect social interactions and alter the behavior of the same-cage littermates of mTBI mice, or alternatively, food-restricted female mice may require more habituation time to express a clear preference for the novel object.

mTBI mice showed consistent impairment in the STARR protocol of the modified Barnes maze, which takes place one week following the initial three days of traditional Barnes maze training. The introduction of the barriers utilized in the STARR protocol led to more inefficient search strategies for the mTBI mice, both male and female. The mTBI group covered significantly more distance than the sham group during the training trials (Figure 7C-D). Notably, mice were tested in the Barnes maze two weeks after mTBI induction, indicating that certain cognitive deficits may last longer than others. The discrepancy between the duration of cognitive deficits found in interval timing as compared to the Barnes maze may reflect differences in assay sensitivity or focal behavioral impairments of mild concussion.

The possible behavioral sex differences following mTBI described here are consistent with those found in other studies that demonstrate that female rodents are less prone to deficits in sensorimotor function, cognition, and anxiety-like behavior following TBI (Rubin & Lipton, 2019). Many have attributed this neuroprotection to female sex hormones such as estrogen and progesterone. Post-injury progesterone and estrogen injections in mice have proved effective at lowering cerebral edema in rats following TBI, highlighting one possible mechanism by which female rodents may be more resistant to post-injury cognitive deficits (Roof et al., 1996; Scott et al., 2022). Other studies on ovariectomized rodents found that both estrogen and progesterone attenuate the increase in blood-brain barrier permeability and cerebral edema following TBI (Kövesdi et al., 2020; O’Connor et al., 2005). Notably, these sex hormones have a myriad of antioxidant and anti-inflammatory properties (Kövesdi et al., 2020). While rodent data on the neuroprotective effects of sex hormones for TBI are robust, human data are limited to observational studies of post-menopausal women undergoing hormone replacement therapy, focusing more so on its effects on cardiovascular disease and cancer rather than on injury and recovery (Gartlehner et al., 2022; Harman, 2006). Nevertheless, rodent models suggest a strong neuroprotective role of estrogen and progesterone for TBIs, which has yet to be fully examined in humans.

In sum, we found evidence that a single mTBI event can cause a range of cognitive and behavioral impairments, particularly in male mice. The male mTBI mice demonstrated spatial working memory errors, interval-timing deficits, and indicators of elevated anxiety-like behavior. Female mTBI mice, on the other hand, were relatively preserved in these cognitive domains with the exception of behavioral flexibility in the STARR protocol of the Barnes maze. Further research will seek to establish the effects of repetitive mTBI events, the protective effects of sex hormones, and whether TBI may cause changes in social behavior.

## TRANSPARENCY, RIGOR, AND REPRODUCIBILITY

All procedures conducted in this study were approved by the Institutional Animal Care and Use Committee (IACUC) at the University of Iowa. 10 male mice and 10 female mice were included in this study, of which 5 of each sex were chosen to be in the mTBI treatment group via random number generator. Behavioral experiments were conducted by researchers blinded to the treatment group. All mice were included in the analysis for each of the behavioral assays. Replication experiments are currently being conducted to establish the consistency of these behavioral effects.

## ACKNOWLEDGMENTS

Funded by a summer fellowship to EBE and JN through the University of Iowa Neurology Department and NINDS NS134833 to GMA. Thank you to Nandakumar Narayanan for contributing his research space and resources for this project.

## AUTHOR CONTRIBUTIONS

JN, AB, GMA, and EBE designed experiments, JN and EBE performed experiments, JN, AB, GMA, and EBE analyzed data, and JN, GMA, and EBE wrote the manuscript.

## References

Acosta, S. A., Tajiri, N., Shinozuka, K., Ishikawa, H., Grimmig, B., Diamond, D., Sanberg, P. R., Bickford, P. C., Kaneko, Y., & Borlongan, C. V. (2013). Long-Term Upregulation of Inflammation and Suppression of Cell Proliferation in the Brain of Adult Rats Exposed to Traumatic Brain Injury Using the Controlled Cortical Impact Model. PLOS ONE, 8(1), e53376. 10.1371/journal.pone.0053376

Antunes, M., & Biala, G. (2012). The novel object recognition memory: Neurobiology, test procedure, and its modifications. Cognitive Processing, 13(2), 93–110. 10.1007/s10339-011-0430-z

Balci, F., Papachristos, E. B., Gallistel, C. R., Brunner, D., Gibson, J., & Shumyatsky, G. P. (2008). Interval timing in genetically modified mice: A simple paradigm. Genes, Brain, and Behavior, 7(3), 373–384. 10.1111/j.1601-183X.2007.00348.x

Bertolli, A., Halhouli, O., Liu-Martínez, Y., Blaine, B., Thangavel, R., Zhang, Q., Emmons, E., Narayanan, N. S., Gumusoglu, S. B., Geerling, J. C., & Aldridge, G. M. (2024). Renovating the Barnes maze for mouse models of Dementia with STARR FIELD: A 4-day protocol that probes learning rate, retention and cognitive flexibility. bioRxiv: The Preprint Server for Biology, 2024.11.30.625516. 10.1101/2024.11.30.625516

Bettis, T., & Jacobs, L. F. (2012). Sex differences in object recognition are modulated by object similarity. Behavioural Brain Research, 233(2), 288–292. 10.1016/j.bbr.2012.04.028

Blaylock, R. L., & Maroon, J. (2011). Immunoexcitotoxicity as a central mechanism in chronic traumatic encephalopathy—A unifying hypothesis. Surgical Neurology International, 2, 107. 10.4103/2152-7806.83391

Boyko, M., Gruenbaum, B. F., Shelef, I., Zvenigorodsky, V., Severynovska, O., Binyamin, Y., Knyazer, B., Frenkel, A., Frank, D., & Zlotnik, A. (2022). Traumatic brain injury-induced submissive behavior in rats: Link to depression and anxiety. Translational Psychiatry, 12(1), 239. 10.1038/s41398-022-01991-1

Diagnostic and statistical manual of mental disorders, fifth edition (DSM-5). (n.d.).

Emmons, E. B., De Corte, B. J., Kim, Y., Parker, K. L., Matell, M. S., & Narayanan, N. S. (2017). Rodent Medial Frontal Control of Temporal Processing in the Dorsomedial Striatum. The Journal of Neuroscience, 37(36), 8718–8733. 10.1523/JNEUROSCI.1376-17.2017

Gardner, R. C., & Yaffe, K. (2015). Epidemiology of mild traumatic brain injury and neurodegenerative disease. Molecular and Cellular Neurosciences, 66(Pt B), 75–80. 10.1016/j.mcn.2015.03.001

Gartlehner, G., Patel, S. V., Reddy, S., Rains, C., Schwimmer, M., & Kahwati, L. (2022). Hormone Therapy for the Primary Prevention of Chronic Conditions in Postmenopausal Persons: Updated Evidence Report and Systematic Review for the US Preventive Services Task Force. JAMA, 328(17), 1747–1765. 10.1001/jama.2022.18324

Gawel, K., Gibula, E., Marszalek-Grabska, M., Filarowska, J., & Kotlinska, J. H. (2019). Assessment of spatial learning and memory in the Barnes maze task in rodents-methodological consideration. Naunyn-Schmiedeberg’s Archives of Pharmacology, 392(1), 1–18. 10.1007/s00210-018-1589-y

Harman, S. M. (2006). Estrogen replacement in menopausal women: Recent and current prospective studies, the WHI and the KEEPS. Gender Medicine, 3(4), 254–269. 10.1016/s1550-8579(06)80214-7

Hernandez-Ontiveros, D. G., Tajiri, N., Acosta, S., Giunta, B., Tan, J., & Borlongan, C. V. (2013). Microglia Activation as a Biomarker for Traumatic Brain Injury. Frontiers in Neurology, 4, 30. 10.3389/fneur.2013.00030

Kane, M. J., Angoa-Pérez, M., Briggs, D. I., Viano, D. C., Kreipke, C. W., & Kuhn, D. M. (2012). A mouse model of human repetitive mild traumatic brain injury. Journal of Neuroscience Methods, 203(1), 41–49. 10.1016/j.jneumeth.2011.09.003

Katz, D. I., Cohen, S. I., & Alexander, M. P. (2015). Mild traumatic brain injury. Handbook of Clinical Neurology, 127, 131–156. 10.1016/B978-0-444-52892-6.00009-X

Kövesdi, E., Szabó-Meleg, E., & Abrahám, I. M. (2020). The Role of Estradiol in Traumatic Brain Injury: Mechanism and Treatment Potential. International Journal of Molecular Sciences, 22(1), 11. 10.3390/ijms22010011

Larson, T., Khandelwal, V., Weber, M. A., Leidinger, M. R., Meyerholz, D. K., Narayanan, N. S., & Zhang, Q. (2022). Mice expressing P301S mutant human tau have deficits in interval timing. Behavioural Brain Research, 432, 113967. 10.1016/j.bbr.2022.113967

Lewon, M., & Hayes, L. J. (2015). The effect of the magnitude of the food deprivation motivating operation on free operant preference in mice. Behavioural Processes, 115, 135–142. 10.1016/j.beproc.2015.03.015

Liu, Y.-W., Zhang, J., Bi, W., Zhou, M., Li, J., Xiong, T., Yang, N., Zhao, L., Chen, X., Zhou, Y., He, W., Yang, T., Wang, H., Xu, L., & Dai, S.-S. (2022). Histones of Neutrophil Extracellular Traps Induce CD11b Expression in Brain Pericytes Via Dectin-1 after Traumatic Brain Injury. Neuroscience Bulletin, 38(10), 1199–1214. 10.1007/s12264-022-00902-0

Lueptow, L. M. (2017). Novel Object Recognition Test for the Investigation of Learning and Memory in Mice. Journal of Visualized Experiments: JoVE, (126), 55718. 10.3791/55718

Luo, J., Nguyen, A., Villeda, S., Zhang, H., Ding, Z., Lindsey, D., Bieri, G., Castellano, J. M., Beaupre, G. S., & Wyss-Coray, T. (2014). Long-Term Cognitive Impairments and Pathological Alterations in a Mouse Model of Repetitive Mild Traumatic Brain Injury. Frontiers in Neurology, 5. 10.3389/fneur.2014.00012

Luo, Y., Zou, H., Wu, Y., Cai, F., Zhang, S., & Song, W. (2017). Mild traumatic brain injury induces memory deficits with alteration of gene expression profile. Scientific Reports, 7(1), 10846. 10.1038/s41598-017-11458-9

Mello, G. B. M., Soares, S., & Paton, J. J. (2015). A Scalable Population Code for Time in the Striatum. Current Biology, 25(9), 1113–1122. 10.1016/j.cub.2015.02.036

Merchant, H., & Averbeck, B. B. (2017). The Computational and Neural Basis of Rhythmic Timing in Medial Premotor Cortex. Journal of Neuroscience, 37(17), 4552–4564. 10.1523/JNEUROSCI.0367-17.2017

Multiple Cause of Death Data on CDC WONDER. (n.d.). Retrieved July 21, 2025, from https://wonder.cdc.gov/mcd.html

Narayanan, N. S., Land, B. B., Solder, J. E., Deisseroth, K., & DiLeone, R. J. (2012). Prefrontal D1 dopamine signaling is required for temporal control. Proceedings of the National Academy of Sciences, 109(50), 20726–20731. 10.1073/pnas.1211258109

O’Connor, C. A., Cernak, I., & Vink, R. (2005). Both estrogen and progesterone attenuate edema formation following diffuse traumatic brain injury in rats. Brain Research, 1062(1–2), 171–174. 10.1016/j.brainres.2005.09.011

Ojo, J.-O., Mouzon, B., Greenberg, M. B., Bachmeier, C., Mullan, M., & Crawford, F. (2013). Repetitive Mild Traumatic Brain Injury Augments Tau Pathology and Glial Activation in Aged hTau Mice. Journal of Neuropathology & Experimental Neurology, 72(2), 137–151. 10.1097/NEN.0b013e3182814cdf

O’Shaughnessy, K. L., Oshiro, W. M., Jackson, T. W., Starnes, H. M., Sasser, A. L., & McMichael, B. D. (2023). Chapter Eight - Neurotoxicity of poly- and perfluoroalkyl substances (PFAS): Epidemiological and rodent studies of behavioral outcomes. In P. R. S. Kodavanti, M. Aschner, & L. G. Costa (Eds.), Advances in Neurotoxicology (Vol. 10, pp. 325–366). Academic Press. 10.1016/bs.ant.2023.09.002

Petraglia, A. L., Plog, B. A., Dayawansa, S., Dashnaw, M. L., Czerniecka, K., Walker, C. T., Chen, M., Hyrien, O., Iliff, J. J., Deane, R., Huang, J. H., & Nedergaard, M. (2014). The pathophysiology underlying repetitive mild traumatic brain injury in a novel mouse model of chronic traumatic encephalopathy. Surgical Neurology International, 5, 184. 10.4103/2152-7806.147566

Pitts, M. W. (2018). Barnes Maze Procedure for Spatial Learning and Memory in Mice. Bio-Protocol, 8(5), e2744. 10.21769/BioProtoc.2744

Roof, R. L., Duvdevani, R., Heyburn, J. W., & Stein, D. G. (1996). Progesterone rapidly decreases brain edema: Treatment delayed up to 24 hours is still effective. Experimental Neurology, 138(2), 246–251. 10.1006/exnr.1996.0063

Rubin, T. G., & Lipton, M. L. (2019). Sex Differences in Animal Models of Traumatic Brain Injury. Journal of Experimental Neuroscience, 13, 1179069519844020. 10.1177/1179069519844020

Scott, M. C., Prabhakara, K. S., Walters, A. J., Olson, S. D., & Cox, C. S. (2022). Determining Sex-Based Differences in Inflammatory Response in an Experimental Traumatic Brain Injury Model. Frontiers in Immunology, 13, 753570. 10.3389/fimmu.2022.753570

Scott, T. L., & Vonder Haar, C. (2019). Frontal brain injury chronically impairs timing behavior in rats. Behavioural Brain Research, 356, 408–414. 10.1016/j.bbr.2018.09.004

Shishido, H., Ueno, M., Sato, K., Matsumura, M., Toyota, Y., Kirino, Y., Tamiya, T., Kawai, N., & Kishimoto, Y. (2019). Traumatic Brain Injury by Weight-Drop Method Causes Transient Amyloid-β Deposition and Acute Cognitive Deficits in Mice. Behavioural Neurology, 2019. 10.1155/2019/3248519

Singh, A., Cole, R. C., Espinoza, A. I., Wessel, J. R., Cavanagh, J. F., & Narayanan, N. S. (2023). Evoked mid-frontal activity predicts cognitive dysfunction in Parkinson’s disease. *Journal of Neurology*, Neurosurgery & Psychiatry, 94(11), 945–953. 10.1136/jnnp-2022-330154

Stern, R. A., Riley, D. O., Daneshvar, D. H., Nowinski, C. J., Cantu, R. C., & McKee, A. C. (2011). Long-term consequences of repetitive brain trauma: Chronic traumatic encephalopathy. *PM & R: The Journal of Injury*, Function, and Rehabilitation, *3*(10 Suppl 2), S460-467. 10.1016/j.pmrj.2011.08.008

Stevanovic, K. D., Fry, S. A., DeFilipp, J. M. S., Wu, N., Bernstein, B. J., & Cushman, J. D. (2022). Assessing the importance of sex in a hippocampus-dependent behavioral test battery in C57BL/6NTac mice. *Learning & Memory (Cold Spring Harbor*, N.Y.), 29(8), 203–215. 10.1101/lm.053599.122

Stutt, H. R., Weber, M. A., Cole, R. C., Bova, A. S., Ding, X., McMurrin, M. S., & Narayanan, N. S. (2024). Sex similarities and dopaminergic differences in interval timing. Behavioral Neuroscience, 138(2), 85–93. 10.1037/bne0000577

Tsao, C.-H., Wu, K.-Y., Su, N. C., Edwards, A., & Huang, G.-J. (2023). The influence of sex difference on behavior and adult hippocampal neurogenesis in C57BL/6 mice. Scientific Reports, 13(1), 17297. 10.1038/s41598-023-44360-8

Vaidya, C. J., & Stollstorff, M. (2008). Cognitive neuroscience of Attention Deficit Hyperactivity Disorder: Current status and working hypotheses. Developmental Disabilities Research Reviews, 14(4), 261–267. 10.1002/ddrr.40

Wang, Y.-S., Hsieh, W., Chung, J.-R., Lan, T.-H., & Wang, Y. (2019). Repetitive mild traumatic brain injury alters diurnal locomotor activity and response to the light change in mice. Scientific Reports, 9(1), 14067. 10.1038/s41598-019-50513-5

Yang, S. H., Gustafson, J., Gangidine, M., Stepien, D., Schuster, R., Pritts, T. A., Goodman, M. D., Remick, D. G., & Lentsch, A. B. (2013). A murine model of mild traumatic brain injury exhibiting cognitive and motor deficits. Journal of Surgical Research, 184(2), 981–988. 10.1016/j.jss.2013.03.075

Yoon, J. H., Minzenberg, M. J., Ursu, S., Walters, R., Wendelken, C., Ragland, J. D., & Carter, C. S. (2008). Association of Dorsolateral Prefrontal Cortex Dysfunction With Disrupted Coordinated Brain Activity in Schizophrenia: Relationship With Impaired Cognition, Behavioral Disorganization, and Global Function. American Journal of Psychiatry, 165(8), 1006–1014. 10.1176/appi.ajp.2008.07060945

